# Deciphering the language of antibodies using self-supervised learning

**DOI:** 10.1101/2021.11.10.468064

**Authors:** Jinwoo Leem, Laura S. Mitchell, James H.R. Farmery, Justin Barton, Jacob D. Galson

## Abstract

An individual’s B cell receptor (BCR) repertoire encodes information about past immune responses, and potential for future disease protection. Deciphering the information stored in BCR sequence datasets will transform our fundamental understanding of disease and enable discovery of novel diagnostics and antibody therapeutics. One of the grand challenges of BCR sequence analysis is the prediction of BCR properties from their amino acid sequence alone. Here we present an antibody-specific language model, AntiBERTa, which provides a contextualised representation of BCR sequences. Following pre-training, we show that AntiBERTa embeddings learn biologically relevant information, generalizable to a range of applications. As a case study, we demonstrate how AntiBERTa can be fine-tuned to predict paratope positions from an antibody sequence, outperforming public tools across multiple metrics. To our knowledge, AntiBERTa is the deepest protein family-specific language model, providing a rich representation of BCRs. AntiBERTa embeddings are primed for multiple downstream tasks and can improve our understanding of the language of antibodies.

## Introduction

B cells are critical to immune protection through their production of antibodies with specific binding properties. To recognize any potential antigen, an individual has a vast diversity of B cells with different B cell receptors (BCRs) – estimated to be as high as 10^15^ variants (1,2). BCR repertoire diversity is generated through the process of somatic recombination of V, D (heavy chain only) and J gene segments during B cell development, followed by somatic hypermutation during B cell activation. Each BCR is comprised of two pairs of heavy-light chain pairs. The heavy chain and light chain each have three complementarity-determining regions (CDRs), which largely determine the BCR’s target specificity. (1,2)

Characterising the BCR repertoire of an individual has proven to be a valuable tool for understanding the fundamental biology of B cells (3) as well as characterising changes during disease (4–7). There are also clinical applications of BCR repertoire analysis in finding novel diagnostics, and therapeutic antibody discovery. Most analyses have focussed on comparing high-level differences in total BCR repertoire metrics between cohorts, such as differences in diversity, number of somatic hypermutations (8), isotype subclass usage, and V(D)J gene segment usage (9,10). To realise the full potential of the data, it will be necessary to understand the specific function of individual BCRs within the context of the entire repertoire.

It has so far proven challenging to predict a BCR’s binding specificity and function from its amino acid sequence alone. Most work focuses on analysing the third CDR (CDR3) of the BCR heavy chain, as it is the greatest determinant of binding; however, predicting CDR3 structure and function is notoriously difficult (11–13). Sequence-dissimilar CDR3s can adopt similar structures (14) and recognise similar regions of a target molecule (15), while small changes in CDR3 sequence can change structure and binding properties (16,17). In addition, BCRs with identical CDR3 sequences but changes elsewhere can have different binding properties (18,19).

One solution is to use representation learning techniques to encode BCR sequences as vectors of real numbers, or “embeddings”. Ideally, the embedding should capture the function of each BCR and contextualise it within the larger BCR universe. In addition, the representations can then be used as inputs for downstream machine learning models (20). Some of the earliest forms of BCR representation learning focussed on calculating physicochemical properties of CDR3 sequences, building k-mer frequency matrices, or constructing position-specific substitution matrices (21–25). Representations from these approaches have previously been used for repertoire classification and antibody structure prediction. While interpretable, these methods depend on hand-crafted features that may miss hidden, or latent, patterns in the data. Furthermore, these methods are context-free; they consider sub-units of the CDR3 sequence as being independent, and do not consider the remainder of the BCR sequence.

Recently, deep learning techniques have shown great promise in learning unobserved patterns from amino acid sequences which relate to their structure and function (26). Neural networks, such as transformers (27,28), are first pre-trained via masked language modelling (MLM) to build a protein language model (LM). General protein LMs such as ProtBERT and ESM-1b can then be used to generate a distributed, contextual representation for each amino acid in a protein sequence. The embeddings from these models then act as a “warm” starting point for various downstream tasks, such as protein structure prediction and protein engineering (26,29,30).

In natural language processing applications, single language models can offer superior performance to their multi-lingual counterparts. Likewise, protein family-specific models are known to outperform general protein models (31–34). Therefore, we advocate for a BCR-specific LM that focusses on the nuances of BCR amino acid sequences.

To date, two LMs have been deployed for BCRs and antibodies: DeepAb (13) and Sapiens (35). DeepAb is a bi-directional long short-term memory (LSTM) network that is pre-trained on 100k paired BCR sequences from the Observed Antibody Space (36,37). As sequence embeddings from DeepAb naturally separate into distinct structural clusters, they can help to produce structural predictions. However, it is unclear if the DeepAb embeddings can be harnessed for tasks beyond antibody structure prediction. Furthermore, LSTMs are typically less performant compared to transformers, in terms of accuracy and speed (27,28). Sapiens is composed of two separate four-layer transformer models that were pre-trained on 20M BCR heavy chains and 19M BCR light chains. Sapiens has been used for antibody humanization and can propose mutations that are near equivalent to those chosen by expert antibody engineers. As with DeepAb, the applicability of Sapiens beyond humanization is unclear. Moreover, most protein LMs use at least 12 transformer layers (26,29,30); by comparison, Sapiens is shallow, and it may not capture the full complexity of BCR sequences.

In this work, we propose AntiBERTa (Antibody-specific Bi-directional Encoder Representation from Transformers), a 12-layer transformer model that is pre-trained on 57M human BCR sequences (42M heavy chains and 15M light chains). We demonstrate that AntiBERTa learns meaningful representations of BCRs, which relate to their B cell origin, activation level, immunogenicity, and structure. We also demonstrate how AntiBERTa can fit in a transfer learning framework by using AntiBERTa representations to predict an antibody’s binding site, the paratope. AntiBERTa improves upon the state-of-art for paratope prediction across multiple metrics, reinforcing the value of a protein-family specific representation learning approach.

## Methods

### Datasets

#### Masked language modelling dataset

To pre-train AntiBERTa, human antibody sequences spanning 61 studies were downloaded from the Observed Antibody Space (OAS) database (36) on 10 March 2021.

Antibody sequences were first filtered out for any sequencing errors, as indicated by OAS. Sequences were also required to have at least 20 residues before the CDR1, and 10 residues following the CDR3. The entire collection of 71.98 million unique sequences (52.89 million unpaired heavy chains and 19.09 million unpaired light chains) was then split into disjoint training, validation, and test sets using an 80%:10%:10% ratio.

In total, the MLM training set comprised 42.3 million heavy chain and 15.3 million light chain sequences, while the MLM validation and MLM test sets each consist of 5.3 million heavy chains and 1.9 million light chains. AntiBERTa is a single model that is trained on both heavy and light chains (Supplementary Figure 1).

#### Paratope prediction dataset

To fine-tune AntiBERTa for paratope prediction, human antibody structural and sequence data were downloaded from SAbDab on 26 Aug 2021 (38). X-ray crystal structures of antibody-antigen complexes binding to proteins or peptides with a resolution of 3.0Å or better were used. Single-chain Fv structures, and structures with missing residues within the IMGT-defined CDRs were omitted. Twelve unorthodox structures were manually removed where the annotated antigen was largely in contact with the framework regions rather than the CDRs (Supplementary Figure 2; Supplementary Table 1).

In total, 1092 redundant heavy chain and 1092 redundant light chain sequences were separately clustered at 99% identity using CD-HIT (39), leading to 462 non-redundant heavy chains and 447 non-redundant light chains.

We then assigned the 909 BCR chains to a V gene cluster by hierarchical clustering (5), and removed BCRs that belonged to a V-gene cluster with fewer than three BCRs (Supplementary Table 2). For example, IGLV7-43 belongs to our ‘VL3’ cluster, which only had two BCRs (PDB: 3T2N, 6WH9); thus, the two BCRs were removed from the set of 909 sequences. In total, six BCR chains were removed using this method. The remaining 903 BCR chains were split into training, validation, and test sets by an 80:10:10 ratio. In total, there are 721 BCR chains in the training set, 91 BCR chains in the validation, and 91 BCR chains in the test set.

### AntiBERTa Pre-Training

The AntiBERTa model was pre-trained using a modified setup of the original RoBERTa-base model (40). The vocabulary is composed of 25 tokens: the standard 20 amino acids and 5 special tokens (<s>, </s>, <pad>, <unk>, <mask>). Each amino acid acts as a token, and no byte-pair encoding was used. The entire BCR sequence is considered as a “sentence”.

AntiBERTa is pre-trained via MLM, which has been used elsewhere (26,29,35). Briefly, 15% of amino acids are chosen for perturbation. Of these, 80% are replaced with the <mask> token, 10% with the original amino acid, and 10% by a random amino acid. The masking ratios have been demonstrated elsewhere to be optimal and we retain these in our work (26,28,41).

During pre-training, the model predicts the original amino acid in the perturbed positions, *M* (Supplementary Figure 3). For a sequence *S* = (*s*_*1*_, *s*_*2*_, … *s*_*l*_) in a batch *B*, the MLM loss is

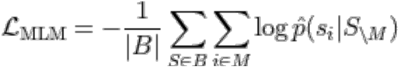

The full set of AntiBERTa hyperparameters is described in Table 1. Briefly, the model was pre-trained for 225000 steps, which equates to three epochs. The learning rate was warmed up to a peak learning rate of 1e-4 over 11250 steps, and linearly decayed thereafter. We used a batch size of 96 across 8 NVIDIA V100 GPUs, for a global batch size of 768.

**Table 1.** Hyperparameters for AntiBERTa, compared to transformers in the wider literature

|  | AntiBERTa | Sapiens | ESM-1b | ProtBERT |
| --- | --- | --- | --- | --- |
| Number of Layers | 12 | 4 | 33 | 30 |
| Number of Attention Heads | 12 | 8 | 20 | 16 |
| Embedding Dimension | 768 | 128 | 1280 | 1024 |
| Feedforward layer dimension | 3072 | 256 | 5120 | 4096 |
| Total number of Parameters | 86M | 569k | 650M | 420M |

A full ablation study of hyperparameters was not performed. However, the 12-layer setup is well-established as a strong baseline for fine-tuning applications (26,30). We envision further hyperparameter sweeps in future work.

### Paratope prediction by AntiBERTa

Paratope prediction was framed as a binary token classification task. For each position in the antibody sequence, AntiBERTa predicts the probability that the position is part of the paratope. This is done by adding a binary classifier “head” on top of AntiBERTa’s 12 layers (Supplementary Figure 4). Paratope prediction was evaluated using six metrics: precision, recall, F1-score, area under the receiver operating characteristic curve (AUC ROC), average precision-recall score (APR), and Matthews’ correlation coefficient (MCC).

Hyperparameters for paratope prediction were estimated by fine-tuning the model over various orders of learning rate magnitude (from 1e-6 to 1e-3), and over different scheduling regimes (constant learning rate, 5% or 10% warmup with linear decay). The optimal setup was decided by the training configuration that yielded the highest APR on the validation set. Experiments were repeated over three seeds to check for variance in the results.

Paratope prediction by AntiBERTa was compared to a PyTorch implementation of Parapred (42); positions with a probability higher than or equal to 0.67 were assigned as the paratope. Sequences were also processed by ProABC-2 (43); since ProABC-2 does not handle unpaired chains, we used paired sequences as input. For example, only the heavy chain of PDB:7KFV is in our test set; for prediction, we submitted both the heavy and light chain sequences of the antibody, but only use the heavy chain predictions for benchmarking. Positions with a ProABC-2 paratope probability higher than or equal to 0.40 were assigned as the paratope.

### BCR representations

For a set of *n* BCR sequences in a batch *B* = (*S*_*1*_, *S*_*2*_, … *S*_*n*_), each with lengths *L* = (*l*_*1*_, *l*_*2*_, … *l*_*n*_), we use the output from the last layer of AntiBERTa. The output embedding is a padded three-dimensional tensor, (*n* x max(*L*) x 768).

For visualisation, we compute the average embedding across the length dimension to generate a two-dimensional (*n* x 768) tensor. Since each batch can comprise different BCR sequence lengths, we omit contributions from padding tokens. As a comparison, we generated BCR embeddings from the last layer of the ProtBERT model, a general protein transformer. Briefly, ProtBERT is a 30-layer transformer model that is trained on a much larger corpus of protein sequences across UniRef and BFD (29).

For testing the representation capabilities of AntiBERTa, two datasets were used: a BCR repertoire dataset containing information on which B cell type each BCR sequence was derived from (44), and 191 non-redundant therapeutic antibodies from TheraSAbDab (45) with anti-drug antibody (ADA) scores. Of these, 65 antibodies also have known source information (46).

## Results

### AntiBERTa learns a meaningful representation of BCR sequences

AntiBERTa is a 12-layer transformer model that is pre-trained on 42 million unpaired heavy chain and 15 million unpaired light chain BCR sequences. Taking inspirations from natural language processing, we consider each BCR sequence as a “sentence”, where each amino acid is a “token”.

AntiBERTa is pre-trained using a self-supervised MLM task, like other transformer-based protein LMs (26,29,35). Briefly, 15% amino acids within the input BCR sequence are randomly perturbed, and the model determines the correct amino acid in place of these masked positions. This task encourages the model to develop a contextual understanding of the BCR sequence. For example, AntiBERTa estimates the probability that an alanine belongs in IMGT 105, given the sequence context (see Methods; Supplementary Figure 3).

Following pre-training, AntiBERTa outputs a distributed vector representation, or embedding, per BCR sequence (see Methods). To visualise the AntiBERTa embeddings, 1000 BCR heavy chains were randomly selected to embed from a well-characterised public dataset (44), then averaged over the length dimension (26,29). Despite only being given BCR sequences and no other information, we find that the BCR embeddings naturally separate according to mutational load and the underlying BCR V gene segments used (Figure 1). Remarkably, there is also good partitioning of BCRs derived from naïve versus memory B cells, suggesting that functionally important information is captured by our model.

**Figure 1.**
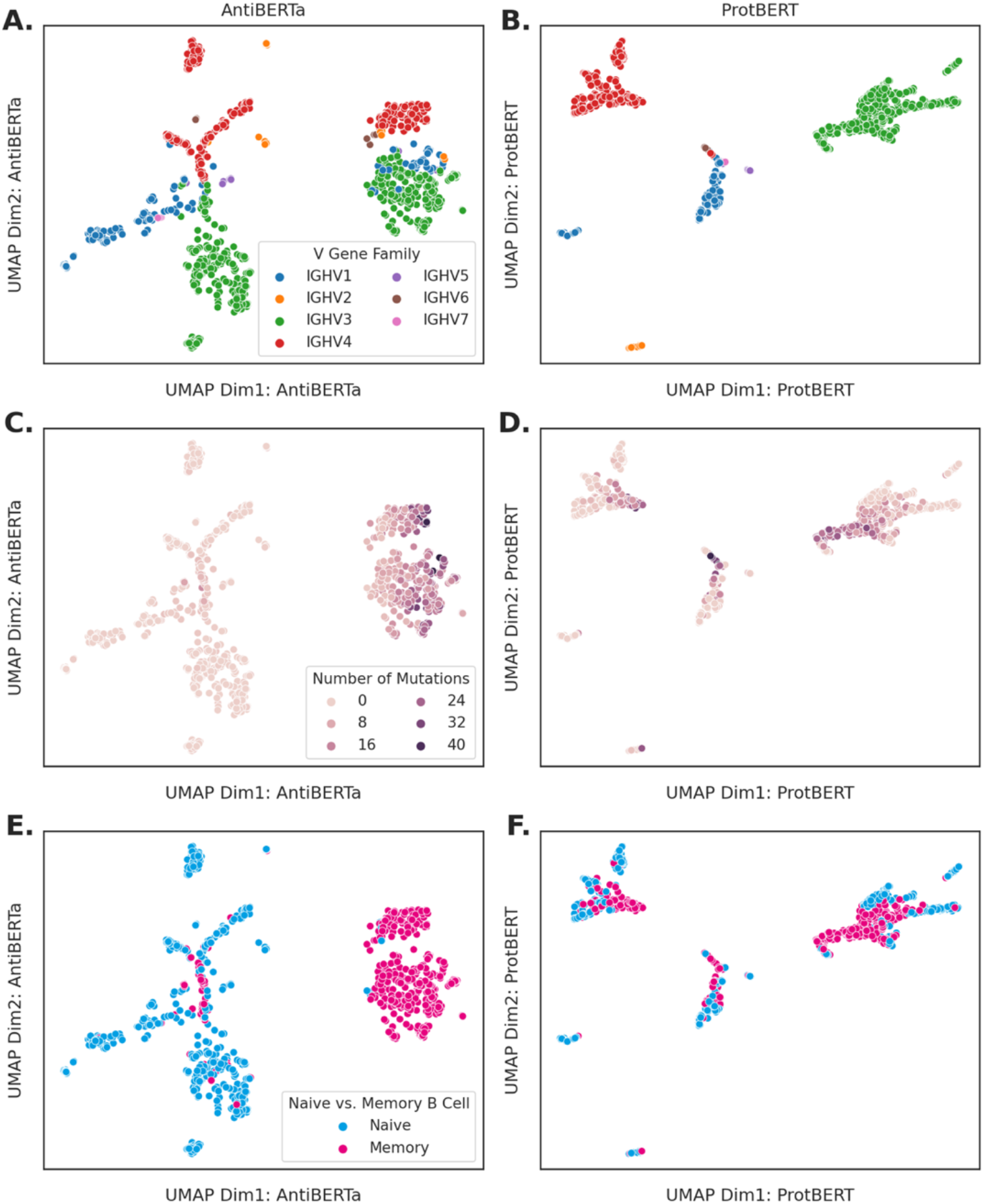
Representation of 1000 sequences randomly selected from (44). Outputs from the final layer of AntiBERTa (A-C) or ProtBERT (D-F) are averaged, then projected onto two dimensions via UMAP (see Methods). Points are coloured according to V gene family (A, D), mutational load (B, E), and B cell population (C, F).

When the same set of BCR sequences are processed via ProtBERT (29), a general protein transformer model, these separations are less distinct. Despite having a smaller dataset and fewer parameters, our BCR-specific transformer captures more patterns that are relevant to BCR function compared to ProtBERT. Furthermore, AntiBERTa has a lower exponentiated cross-entropy for the MLM task (AntiBERTa ECE = 1.43; ProtBERT ECE = 1.72), suggesting that AntiBERTa produces higher-quality representations of BCRs.

We also used AntiBERTa to embed the heavy chains from 198 well-characterised therapeutic antibodies (46,47). AntiBERTa is generally able to separate therapeutic antibodies according to their origin (i.e., chimeric, humanized, human or murine), which also coincides with their identity to their closest human germline V gene (Figure 2). These antibodies also have known anti-drug antibody (ADA) response scores, and we see the separations in ADA largely corresponded to separations by human germline V gene identity. The embeddings therefore offer a method to filter antibodies with potentially high ADA scores and discover safer therapeutics.

**Figure 2.**
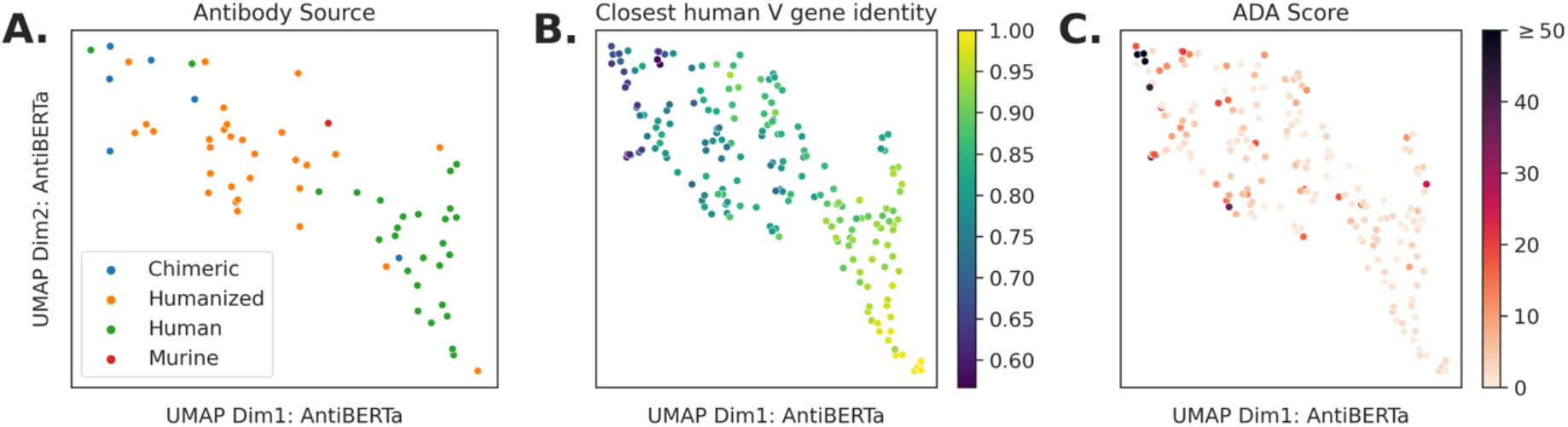
Embedding of 191 non-redundant therapeutic antibodies. Embeddings from the last layer of AntiBERTa are averaged over the length dimension and projected onto two dimensions via UMAP. Each point represents a BCR, and is coloured by antibody source (A), germline V gene identity (B), and ADA score (C).

### Self-attention can provide clues on structure and function

One of the major components of transformer-based models, like AntiBERTa, is its multi-head self-attention mechanism (27). AntiBERTa’s 12 attention heads in each of its 12 layers focus on different aspects of the BCR sequence (Figure 3, Supplementary Figure 5, Supplementary Figure 6). The self-attention scores are then used to compute the final, contextual embedding for each amino acid in the BCR sequence. Typically, self-attention in AntiBERTa tends to be directed toward non-germline positions of the BCR sequence, or between CDR3 positions.

**Figure 3.**
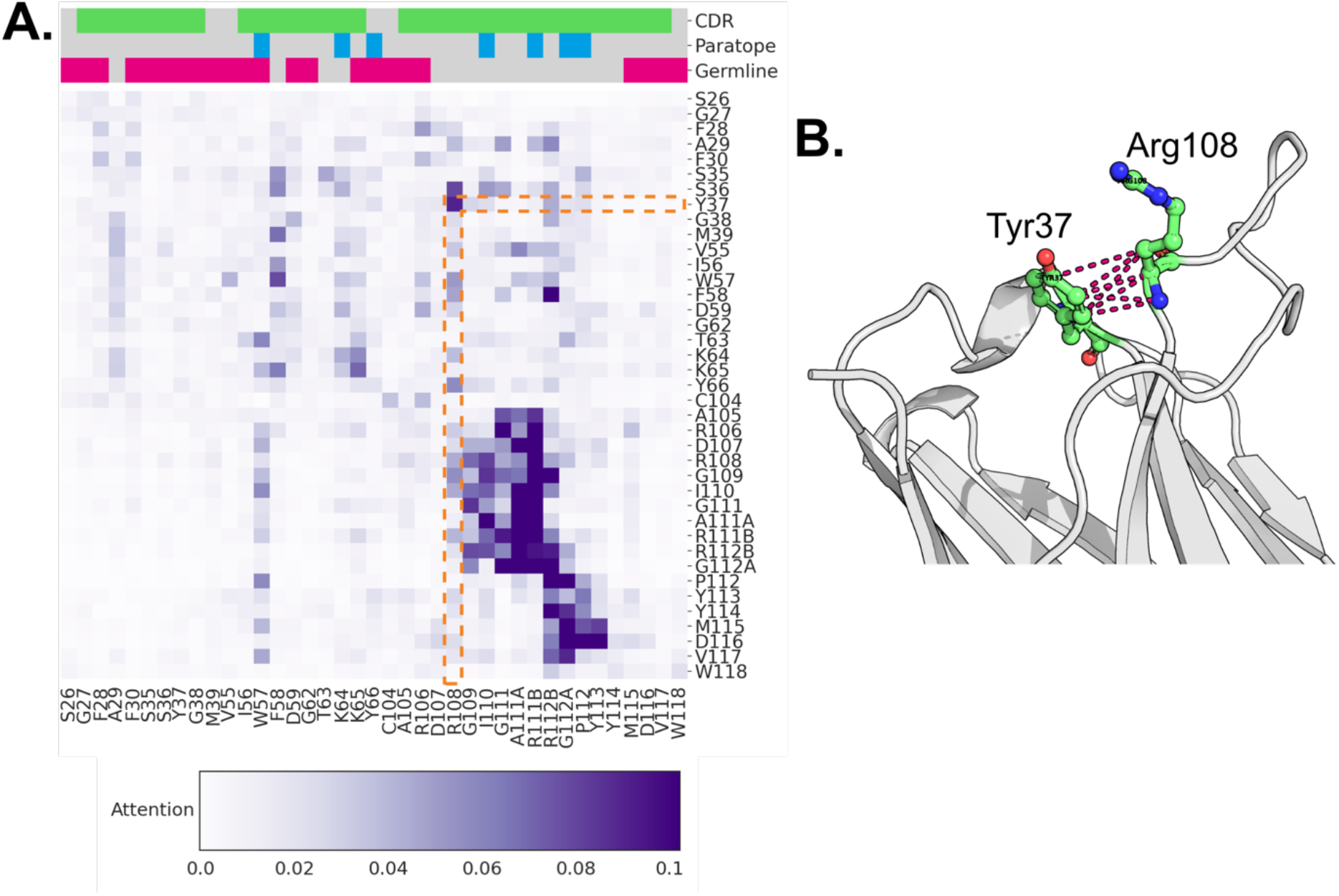
Self-attention heatmap from AntiBERTa’s 12^th^ layer, 6^th^ head, for aducanumab’s heavy chain can show potential structural contacts. A) Attention is directed from positions in the rows toward positions in the columns; stronger self-attention is indicated by darker shades of purple. For each position, we label which positions are germline (pink), part of the paratope (blue), or the CDR (green). Here, we show attention between positions in the CDRs and one position before and after each CDR; the full self-attention map is shown in Supplementary Figure 5. We highlight Tyr37 and Arg108 in orange. B) This was later confirmed from the crystal structure of aducanumab to be a contact (PDB: 6CO3).

We find that residue pairs with high self-attention scores can reveal long-range structural contacts. As an example, we embedded the heavy chain of aducanumab, a recently approved therapeutic antibody binding beta-amyloid. The sixth attention head in AntiBERTa’s final layer places high self-attention between Tyr37 of CDR1 and Arg108 of CDR3 (Figure 3, Supplementary Figure 5). These positions were later confirmed to be a contact within the crystal structure (PDB: 6CO3). Self-attention may also give clues toward functionally interesting antibody positions; the germline residue Trp57 in Aducanumab receives a high level of attention from other residues within the heavy chain (Supplementary Figure 6). This position was then verified to be part of the paratope (48).

When aducanumab is processed by ProtBERT, the self-attention pattern does not show as strong a relationship to non-germline or potential paratope positions (Supplementary Figure 7). We found that ProtBERT pays attention to the disulphide bridge between Cys23-Cys104 (36), while AntiBERTa does not. This further reflects how AntiBERTa pays more attention to what is functionally important for specific binding, as the cysteine pair is almost always invariant for all antibodies.

### Paratope Prediction using AntiBERTa

Pre-trained LMs provide useful representations for transfer learning on a wide range of tasks (28). For example, in natural language processing, pre-trained word embeddings from BERT are fine-tuned for classifying sentences, computing sentence entailment, and named-entity recognition. Similarly, protein representations from LMs such as ESM-1b and ProtBERT have been used for secondary structure prediction and contact prediction (26,29).

As AntiBERTa’s representations seemed to capture hints of BCR function, and self-attention was concentrated on putative paratope positions, we fine-tuned the model for paratope prediction, akin to named-entity recognition. A rapid, accurate, paratope prediction method can shed light on binding properties, and is valuable for therapeutic antibody engineering (42,43,49). While we focus on paratope prediction here, the AntiBERTa model can potentially be fine-tuned for other tasks such as antibody structure prediction and humanisation (13,35).

For each position in the antibody sequence, we predict the probability that it is part of the antibody’s paratope (see Methods). To evaluate paratope prediction, we report the precision, recall, F1-score, MCC, AUROC and APR scores on a held-out test set of 91 antibodies (Table 2). We also benchmark AntiBERTa against two other publicly available tools: Parapred, and ProABC-2 (see Methods). Parapred uses an LSTM and convolutional neural networks (CNNs) to predict paratope positions of the Chothia-defined CDR loops, while ProABC-2 uses CNNs on paired, full-length sequences to predict paratopes.

**Table 2.** Performance metrics of paratope prediction

|  | Precision | Recall | F1 | MCC | AUROC | APR |
| --- | --- | --- | --- | --- | --- | --- |
| Parapred | 0.646 | **0.741** | **0.683** | 0.564 | 0.888 | 0.717 |
| ProABC-2 | 0.662 | 0.584 | 0.621 | 0.582 | 0.951 | 0.655 |
| AntiBERTa | **0.743** | 0.625 | 0.679 | **0.649** | **0.963** | **0.763** |

AntiBERTa predicts paratopes of both CDR and non-CDR positions, like ProABC-2. For the C1A-B12 antibody (PDB: 7KFV; (50)), a SARS-CoV2 binder in our test set, AntiBERTa detects Tyr66 in the framework as a paratope position (Figure 4A, B). Likewise, for the light chain of tralukinomab (PDB: 5L6Y; (51)), another example in our test set, AntiBERTa correctly predicts paratope positions outside of the CDRs (Figure 4C, D). AntiBERTa’s self-attention changes via fine-tuning, suggesting that it adapts its self-attention toward predicting paratope positions. For instance, some CDR1 positions’ self-attention scores for the C12A-B12 antibody strengthen after fine-tuning, while some framework positions’ attentions are redistributed to the CDR3 (Supplementary Figure 8).

**Figure 4.**
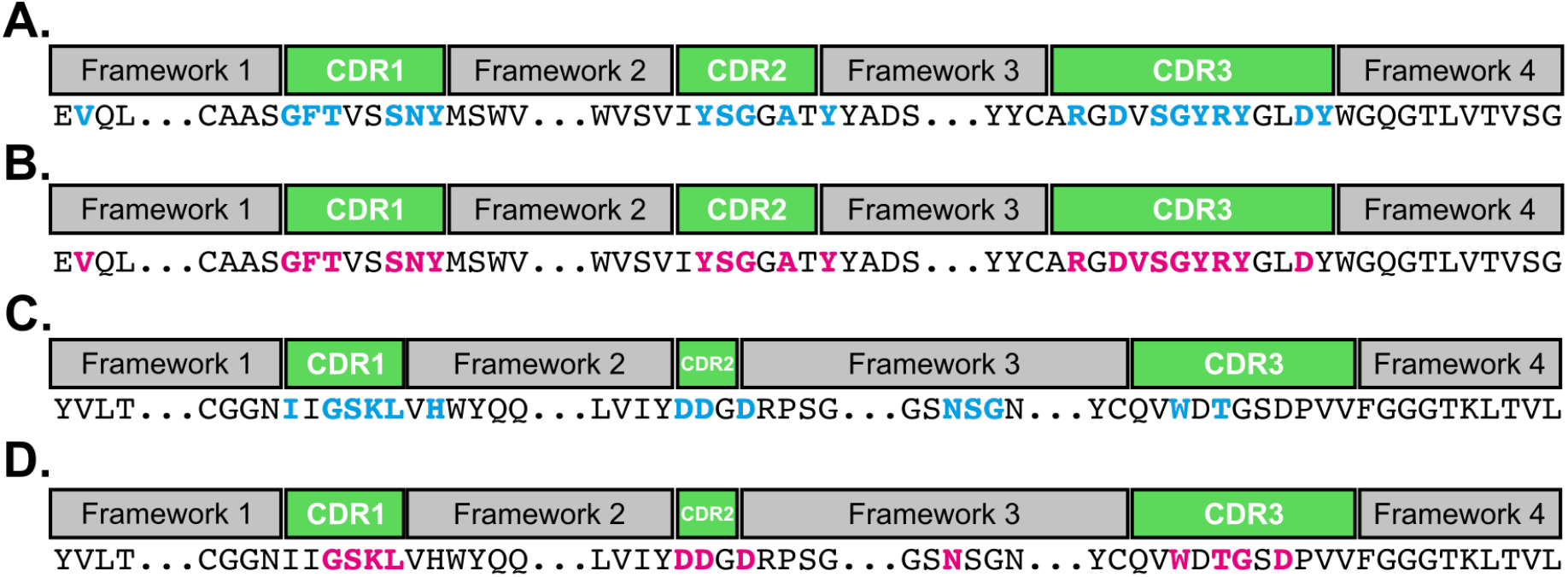
AntiBERTa can predict non-CDR positions that form the paratope. The framework and CDR regions of the C1A-B12 antibody heavy chain (A, B) and tralokinumab light chain (C, D) are outlined in grey and green boxes. A, C) Observed paratope positions from the crystal structure are highlighted with blue letters. B, D) Predicted paratope positions from AntiBERTa are highlighted in pink.

**Figure 5.**
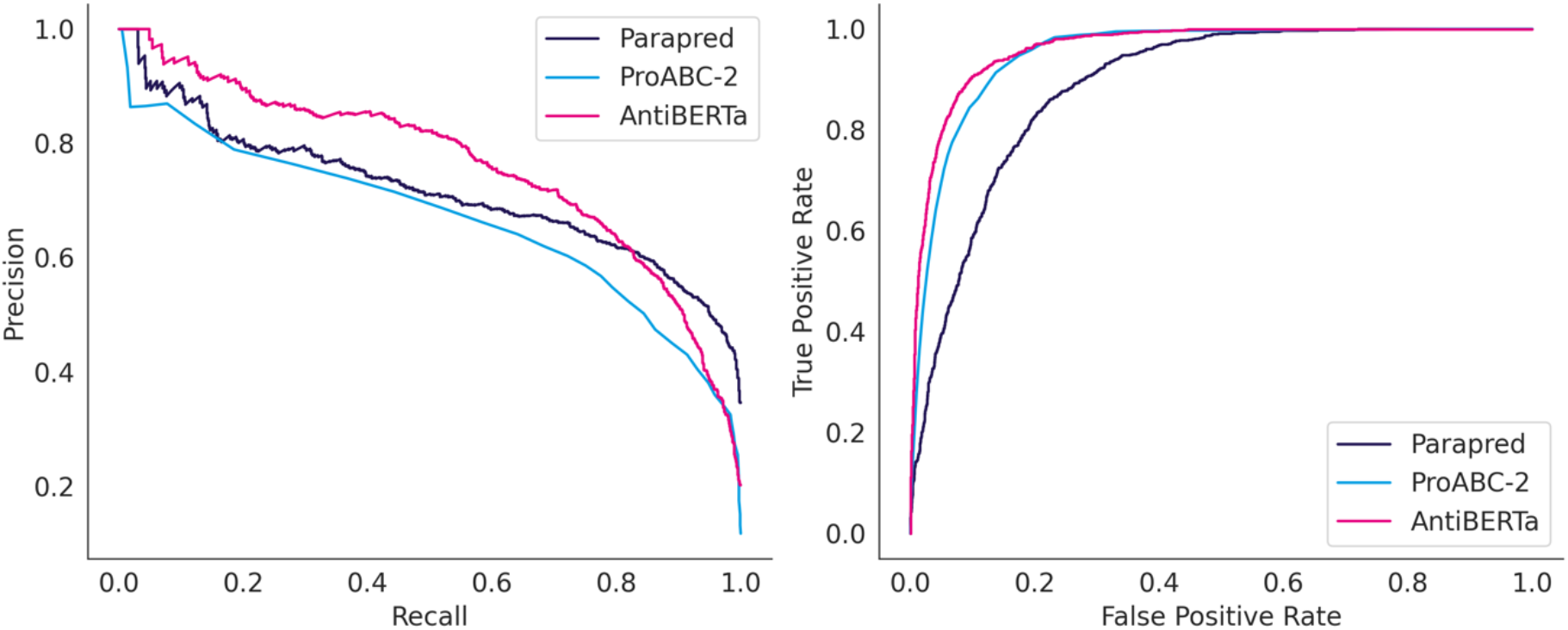
Precision-recall (A) and ROC (B) curves for paratope prediction by Parapred, ProABC-2, and AntiBERTa.

Our transformer-based model is superior to Parapred and ProABC-2 across multiple metrics (Table 2). AntiBERTa has the highest precision, MCC, AUROC, and APR, while Parapred has the highest recall and F1 score. However, Parapred has the lowest precision, which suggests that it is potentially susceptible to a high false positive rate.

When breaking down the predictions by V gene segment family, we find that precision is largely related to the amount of structural data available (Supplementary Figure 9). Precision and recall are unaffected by antigen class, despite the training set having five-fold more protein binders than peptide binders (Supplementary Figure 10).

## Discussion

Here we present AntiBERTa, a transformer-based BCR-specific LM that learns a meaningful representation of BCR sequences. We demonstrate that embeddings from AntiBERTa reflect various biologically meaningful aspects of a BCR, such as mutation count, V gene provenance, B cell origin, and immunogenicity, despite not having this information during pre-training. A key driver of AntiBERTa’s understanding is its multi-head self-attention mechanism, which focusses on structurally and functionally important residues within a BCR sequence. Given these capabilities, we fine-tune the model for paratope prediction to demonstrate the quality of the representations and find that AntiBERTa is the best performer across multiple metrics.

Machine learning methods have previously been used to classify B cells based on the BCR sequence and its features, such as the CDR3 region’s physicochemical properties (44). While we have not explicitly predicted B cell subsets using AntiBERTa, its embeddings can already separate naïve and memory B cells. This shows the advantages of a transformer approach: the model learns latent features of BCRs that correspond to various facets of BCR function, such as its B cell origin. The onus of hand-crafting features that best correlate with BCR origin is effectively delegated to the training process.

A particular benefit of using transformer-based methods is the availability of self-attention heatmaps, which can help explain what the model understands about BCRs. In general, AntiBERTa’s self-attention focusses on non-germline positions. We also find that self-attention can hint toward pairs of residues that contact each other or identify putative paratope positions. While the current self-attention scores do not always carry a clear relationship to antibody structure and function, the self-attention scores may point to latent features that are not yet directly relatable to our current intuitions. A more in-depth analysis of how specific layers and heads correlate with antibody structure and function, as has been done for general proteins, could be the first step in facilitating interpretation (52). Furthermore, some of the self-attention can be noisy; we expect the scores to have sharper focus with more data and a longer pre-training regime.

AntiBERTa outperformed current paratope prediction approaches across various metrics, and its performance is fuelled by a shift in its self-attention scores. One advantage of AntiBERTa is its ability to predict paratope positions outside of the CDRs, meaning it can be used to inform the engineering of non-CDR positions. Furthermore, AntiBERTa can predict the paratopes of unpaired chains, making it suitable for most repertoire datasets where only the heavy or light chain sequences are available. In our current analyses, we did not apply any thresholds on the predicted paratope probabilities, though we envision more work in this space to achieve a more precise predictor (27,38).

Throughout this work, we have visualised BCR sequences as the averaged embedding across the length of the BCR, akin to ProtBert and ESM-1b (26,29). Though there are several strategies to represent full-length sequences (53), we have not explored these in extensive detail here. The optimal method of embedding full-length BCRs may depend on the use case of interest, and we see this as an active area of research in the future.

AntiBERTa offers a high-quality representation of BCR sequences that captures aspects of a BCR’s origin, structure, and function. The embeddings from AntiBERTa also provide a representation of BCRs that can be leveraged for various downstream tasks via a transfer learning paradigm. Specifically, we show that AntiBERTa representations can fuel paratope prediction capabilities. Beyond paratope prediction, we see AntiBERTa being able to empower a wider range of tasks relating to BCR repertoire analysis.

