## Supplementary Information for "Deciphering the language of antibodies using self-supervised learning"

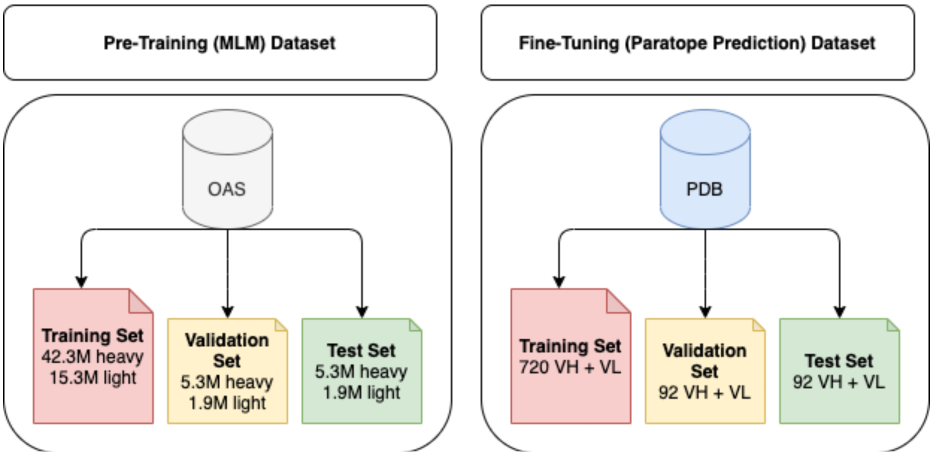


**Supplementary Figure 1**. Datasets used for pre-training and fine-tuning AntiBERTa.

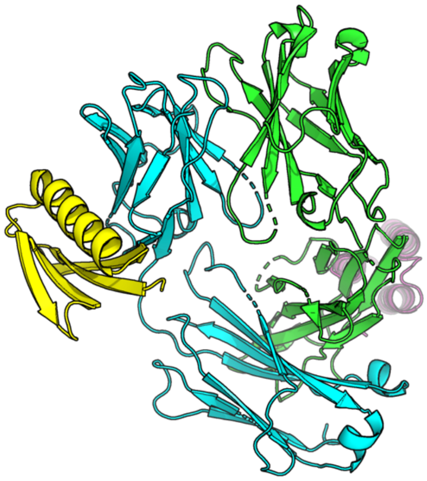

**Supplementary Figure 2.** Example structure that is omitted from analyses, as the antigen is in an unorthodox position with respect to the antibody. PDB: 5U3D.

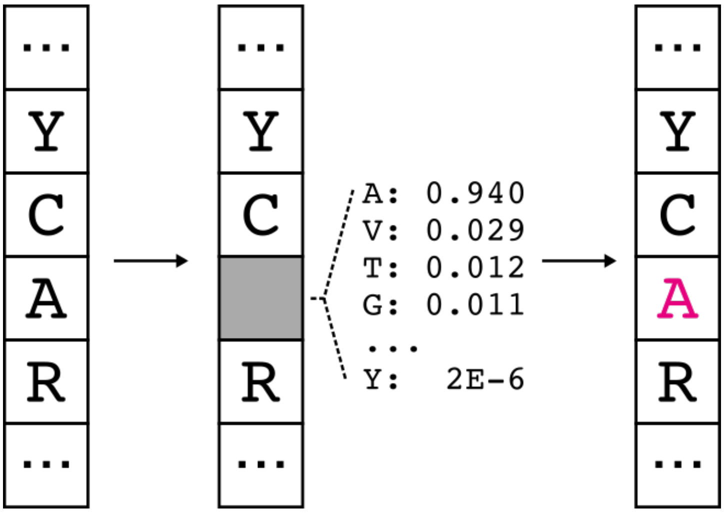


**Supplementary Figure 3.** Masked language modelling (MLM) task for AntiBERTa pre-training. For each BCR sequence, 15% of its positions are perturbed as a mask (grey square), an incorrect amino acid, or the original amino acid (both not shown). For the masked positions, AntiBERTa learns the probability distribution of the correct token given the sequence context, and fills it back in.

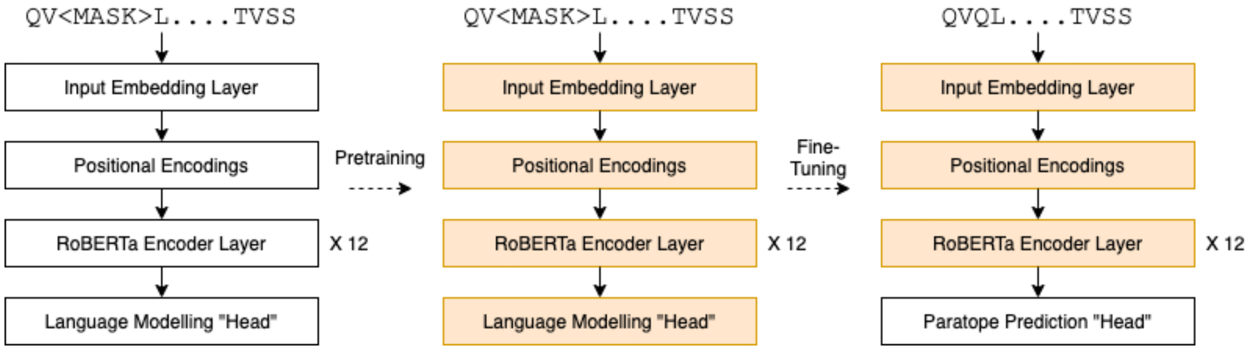


**Supplementary Figure 4.** Model architecture for paratope prediction. AntiBERTa is a 12-layer transformer based on the RoBERTa-base model. After initial pre-training on the MLM task, the model is fine-tuned for paratope prediction. The learned weights of every layer from the MLM task, except for the last head, are used to initialise the paratope predictor model.

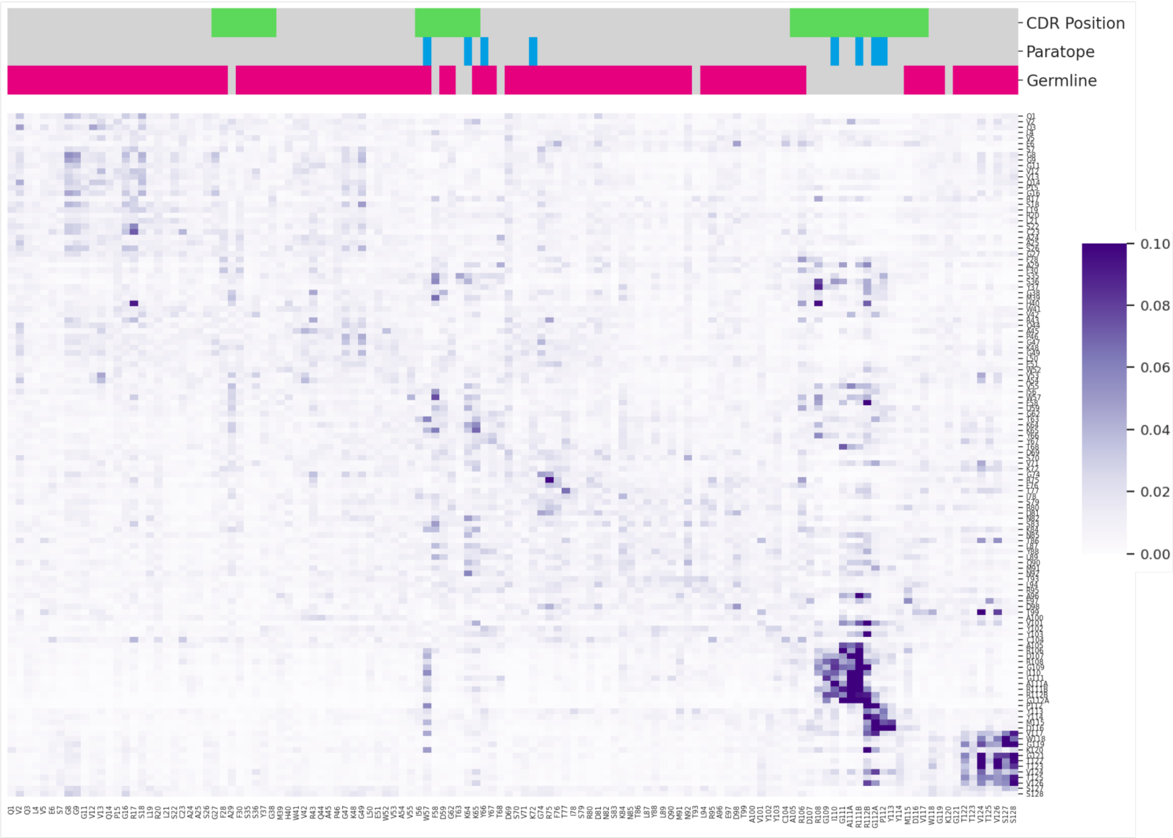


**Supplementary Figure 5.** Self-attention heatmap from AntiBERTa's 12th layer, 6th head for the full sequence of aducanumab's heavy chain.

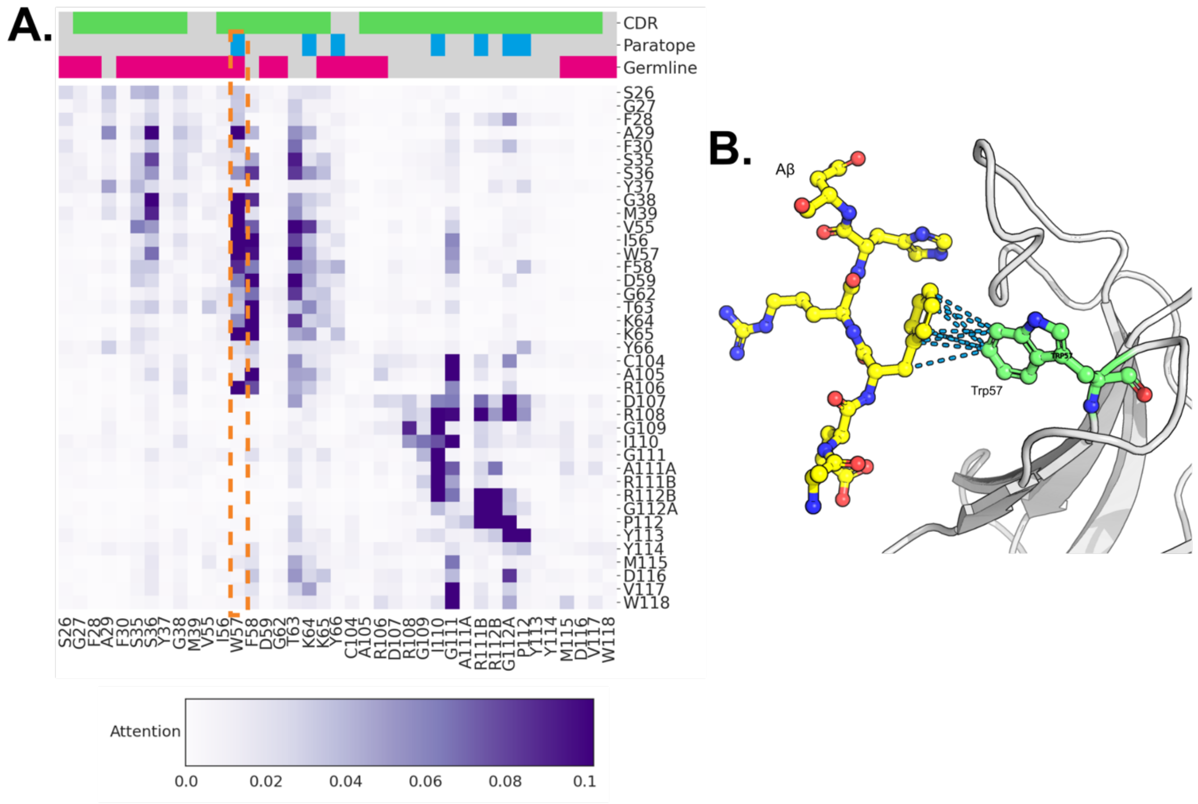


**Supplementary Figure 6.** Self-attention heatmap from AntiBERTa for Aducanumab for layer 12, head 2. Only the CDRs and 1 position before and after each CDR are visualised. The self-attention scores (A) are high for some germline positions, which were later confirmed to be part of the paratope, such as Tyr57 (B).

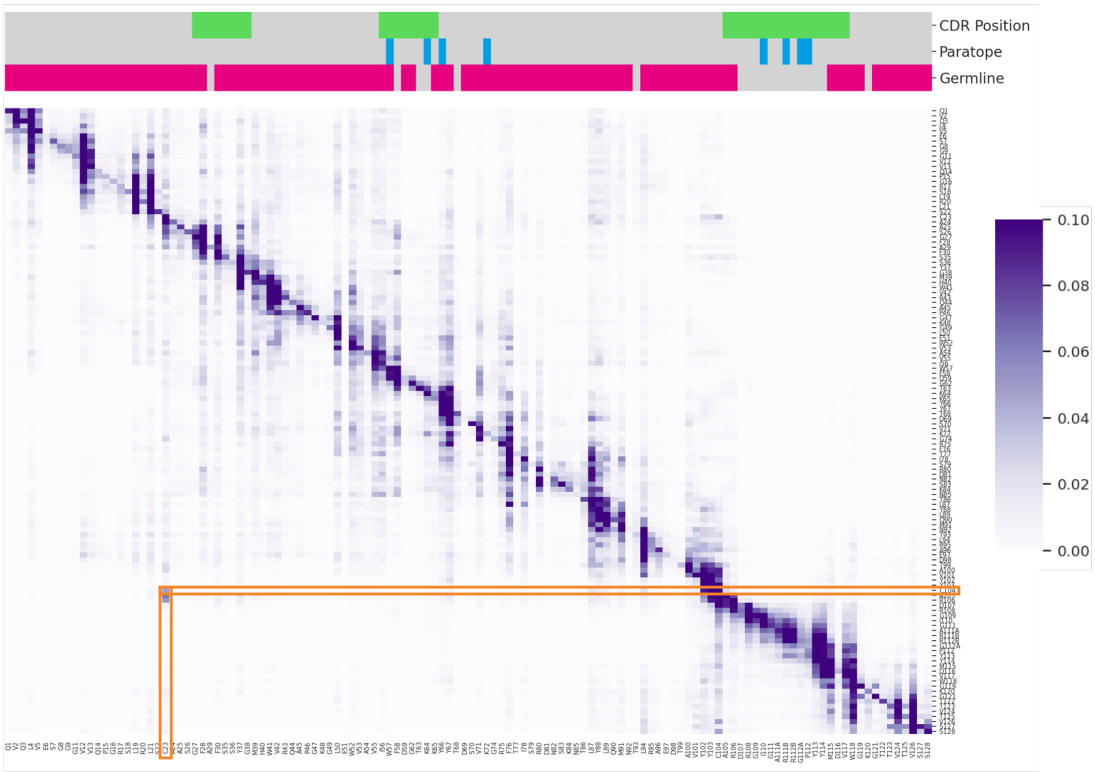


**Supplementary Figure 7.** Self-attention heatmap from ProtBERT layer 29, head 6, for the full sequence of aducanumab's heavy chain. Self-attention between Cys104 to Cys23 is outlined in orange.

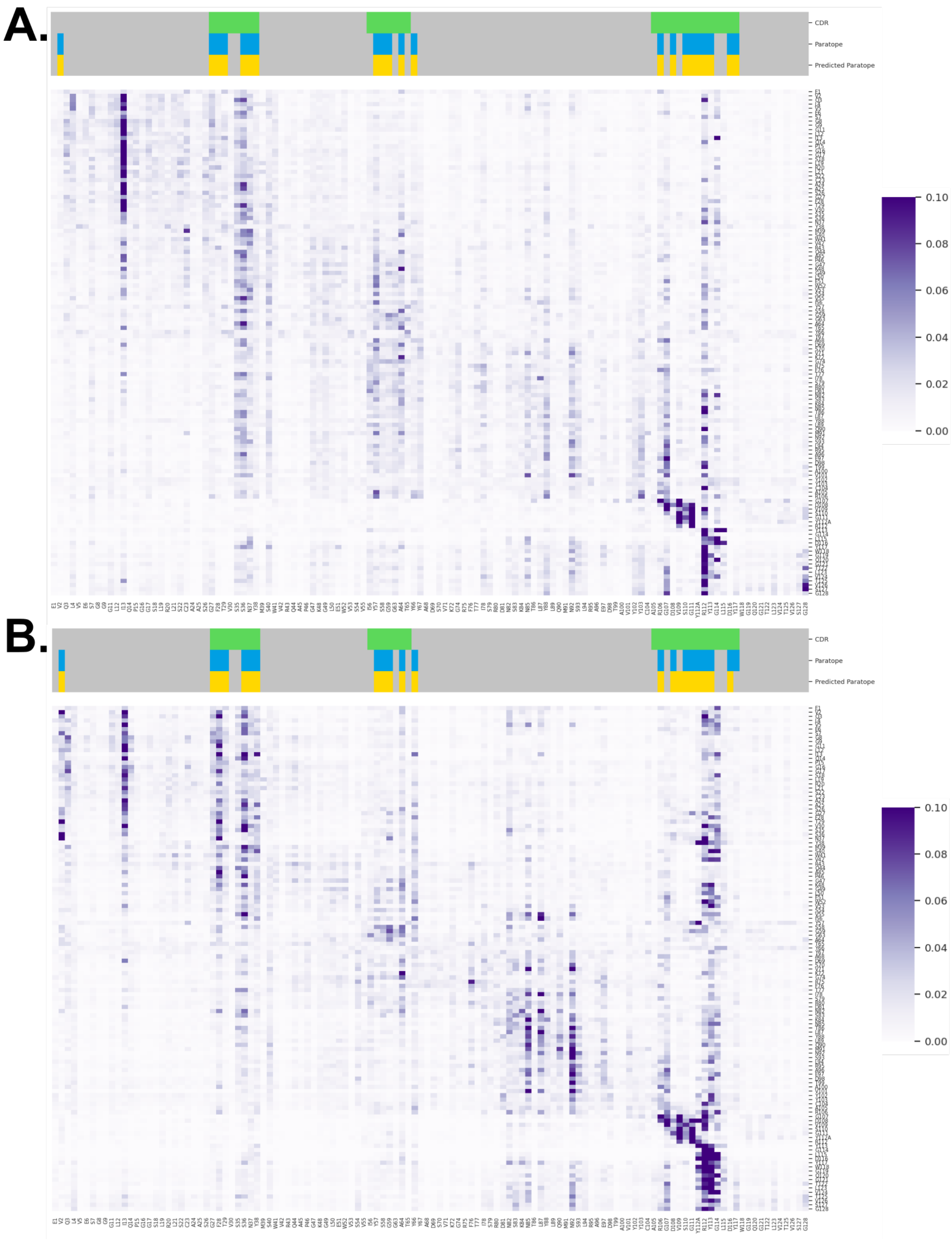


**Supplementary Figure 8.** Self-attention maps from the second attention head of AntiBERTa's 12th layer, before (A) and after (B) fine-tuning. Example shown for the heavy chain of C1A-B12 (PDB: 7KFV), a SARS-CoV2 binding antibody in our test set. We computed its self-attention using AntiBERTa before and after fine-tuning. AntiBERTa pays attention to some paratope positions in the CDR1, which is strengthened after fine-tuning. Some of the attention scores that were distributed throughout framework 1 are redistributed to CDR3 positions after fine-tuning.

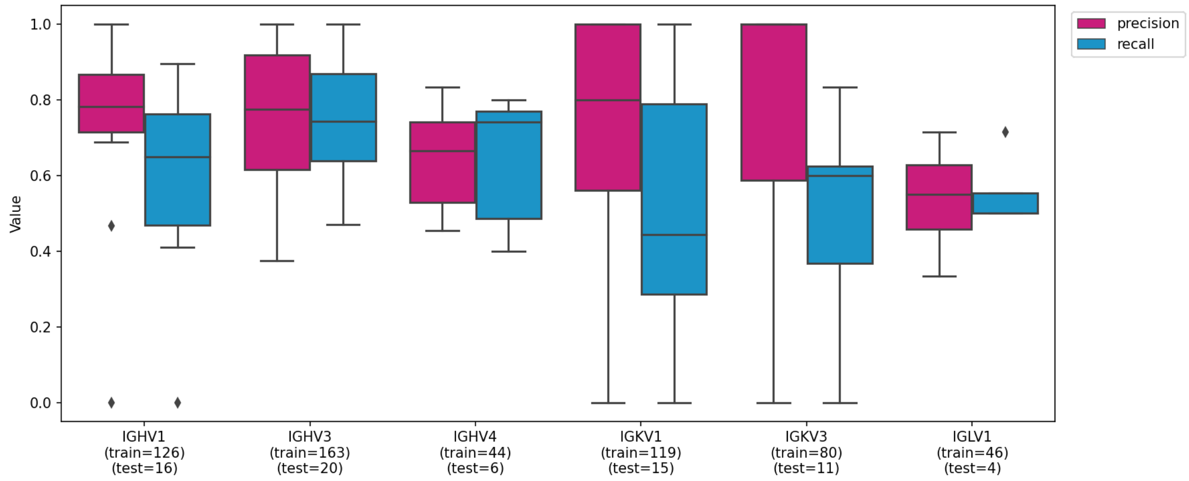


**Supplementary Figure 9.** Breakdown of paratope prediction performance on the test set by germline V gene family. Only V gene families with at least 5 examples in the test set are shown.

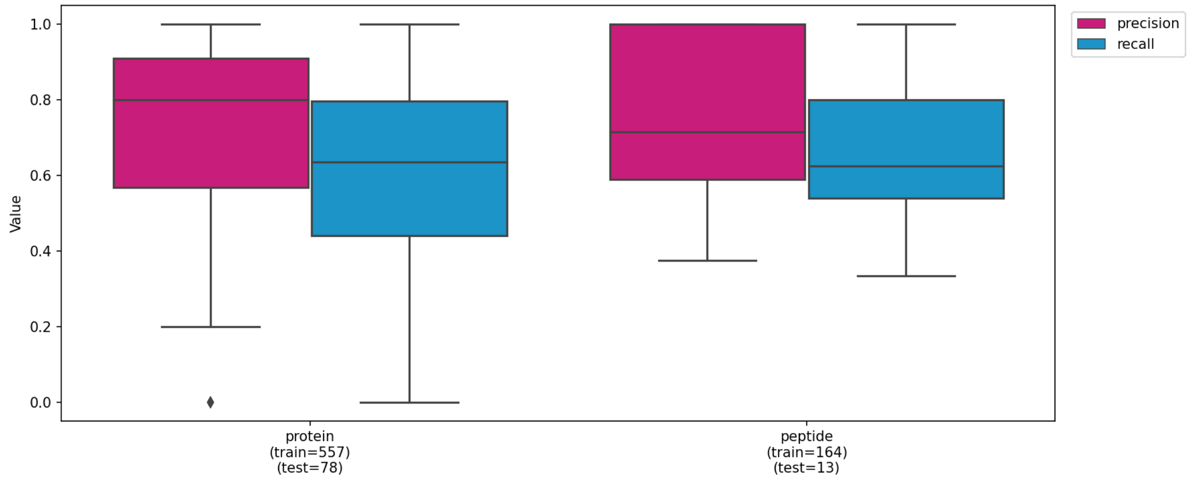


**Supplementary Figure 10.** Breakdown of paratope prediction performance on the test set by antigen class.

**Supplementary Table 1**. List of structures manually removed from analyses due to non-canonical contacts with their annotated antigen.

| | | | |
|---|---|---|---|
| **4ERS** | **4IOI** | **4HJG** | **6W7S** |
| **5U5F** | **5U5M** | **5U6A** | **6B9Y** |
| **6B9Z** | **6BAE** | **6BAH** | **5U3D** |

**Supplementary Table 2.** V-gene clusters used for partitioning the fine-tuning training, validation, test sets.

| Cluster | V genes | Removed PDB entries |
|---|---|---|
| VH1 | IGHV2-5, IGHV2-26, IGHV2-70 | |
| VH2 | IGHV4-4, IGHV4-30-4, IGHV4-31, IGHV4-34, IGHV4-38-2, IGHV4-39, IGHV4-59, IGHV4-61 | |
| VH3 | IGHV6-1 | |
| VH4 | IGHV3-15, IGHV3-49, IGHV3-72, IGHV3-73 | |
| VH5 | IGHV3-7, IGHV3-9, IGHV3-11, IGHV3-13, IGHV3-20, IGHV3-21, IGHV3-23, IGHV3-30, IGHV3-30-3, IGHV3-33, IGHV3-48, IGHV3-53,IGHV3-64,IGHV3-66,IGHV3-74, IGHV3-NL1 | |
| VH6 | IGHV5-10-1, IGHV5-51 | |
| VH7 | IGHV1-24, IGHV1-69-2 | |
| VH8 | IGHV1-2, IGHV1-3, IGHV1-8, IGHV1-18, IGHV1-46, IGHV1-69 | |
| VH9 | IGHV1-58 | 6MG7 |
| VH10 | IGHV7-4-1 | |
| VK1 | IGKV2-28, IGKV2-30 | |
| VK2 | IGKV6-21 | 4G6J |
| VK3 | IGKV1-5, IGKV1-9, IGKV1-12, IGKV1-13, IGKV1-16 IGKV1D-16, IGKV1-17, IGKV1-27, IGKV1-33, IGKV1-39, IGKV1-NL1 | |
| VK4 | IGKV3D-7, IGKV3-11, IGKV3-15, IGKV3-20, IGKV3D-20, IGKV4-1 | |
| VL1 | IGLV4-69 | 5IFJ, 7CR5 |
| VL2 | IGLV5-37, IGLV5-45 | |
| VL3 | IGLV7-43, IGLV7-46 | 3T2N, 6WH9 |
| VL4 | IGLV1-40, IGLV1-44, IGLV2-14, | |
